# Increasing seed thiamin content impacts stored carbon partitioning and subsequent seedling stress tolerance

**DOI:** 10.1101/2021.08.03.454933

**Authors:** Mohammad Yazdani, James A Davis, Jeffrey F Harper, David K Shintani

**Author notes:** **Author for contact:** David K Shintani. **One sentence summary:** Overexpression of thiamin biosynthetic genes in seeds increases seed thiamin level, alters seed carbon partitioning, and increases seedling stress tolerance. **Authors’ contributions:** DKS, JFH, and MY conceived and designed the experiments. MY performed all experiments. JAD aided with lipid analysis by GC. MY and DKS analyzed the results and wrote the manuscript. All authors contributed to the final manuscript. The author responsible for distribution of materials integral to the findings presented in this article is: David K Shintani.

## Abstract

Thiamin and thiamin pyrophosphate (TPP) are essential components for the function of enzymes involved in the metabolism of carbohydrates and amino acids in living organisms. In addition to its role as a cofactor, thiamin plays a key role in resistance against biotic and abiotic stresses in plants. Most of the studies used exogenous thiamin to enhance stress tolerance in plants. In this study, we achieved this objective by genetically engineering *Arabidopsis thaliana* and *Camelina sativa* for the seed-specific co-overexpression of the *Arabidopsis* thiamin biosynthetic genes *Thi4*, *ThiC*, and *ThiE*. Elevated thiamin content in the seeds of transgenic plants was accompanied by the enhanced expression levels of transcripts encoding thiamin cofactor-dependent enzymes. Furthermore, seed germination and root growth in thiamin over-producing lines were more tolerant to oxidative stress caused by salt and paraquat treatments. The transgenic seeds also accumulated more oil (up to16.4% in *Arabidopsis* and17.9% in *C. sativa*) and carbohydrate but less protein than the control seeds. The same results were also observed in TPP over-producing *Arabidopsis* plants generated by the seed-specific overexpression of *TPK1*. Together, our findings suggest that thiamin and TPP over-production in transgenic lines confer a boosted abiotic stress tolerance and alter the seed carbon partitioning as well.

## INTRODUCTION

Thiamin (vitamin B_1_) in its active form TPP is an essential cofactor for the function of key enzymes such as transketolase (TK), pyruvate dehydrogenase (PDH), and α-ketoglutarate dehydrogenase (α-KGDH) which are involved in central carbon metabolism in living organisms (Jordan, 2003; Nosaka, 2006; Goyer, 2010). Humans and animals can convert free thiamin to TPP, but they are unable to synthesize free thiamin *de novo*. Hence, they must take it up from their food to maintain a normal metabolism (Roje, 2007; Kowalska and Kozik, 2008; Goyer, 2010). However, important crop plants including wheat, maize, and rice are not good sources of thiamin (Fitzpatrick et al., 2012), and thiamin deficiency is widespread among people whose diet is largely based on these low-thiamin staple crops (Lonsdale, 2006; Dong et al., 2015). Severe vitamin B_1_ deficiency causes the lethal disease beriberi in humans (Lonsdale, 2006; Roje, 2007; Minhas et al., 2018). Most of the key enzymes involved in thiamin *de novo* biosynthesis and salvage pathways have been identified in plants and bacteria (Jurgenson et al., 2009; Goyer, 2010; Yazdani et al., 2013; Zallot et al., 2014). In plants, thiamin is synthesized by the condensation of 4-amino-5-hydroxymethyl-2-methylpyrimidine pyrophosphate (HMP-PP) and 4-methyl-5-(β*-*hydroxyethyl)thiazole phosphate (HET-P) and is mediated by a bifunctional enzyme known as thiamin-phosphate pyrophosphorylase or ThiE (Fig. 1). These thiamin precursors are biosynthesized through two independent pathways mediated by thiazolephosphate synthase (Thi4) and phosphomethylpyrimidine synthase (ThiC) enzymes (Roje, 2007; Goyer, 2010, 2017). Thi4 has also been shown to be a single turnover or suicide enzyme (Chatterjee et al., 2011). At the end of the pathway (Fig. 1), free thiamin is subjected to pyrophosphorylation by thiamin pyrophosphokinases enzymes (TPK1 and TPK2) to produce its active form TPP (Ajjawi et al., 2007a). In addition to *de novo* biosynthesis, plants have salvage pathways in which the thiamin degradation products can be recycled by two different enzymes known as ThiM and TenA_E (Yazdani et al., 2013; Zallot et al., 2014).

**Figure 1:**
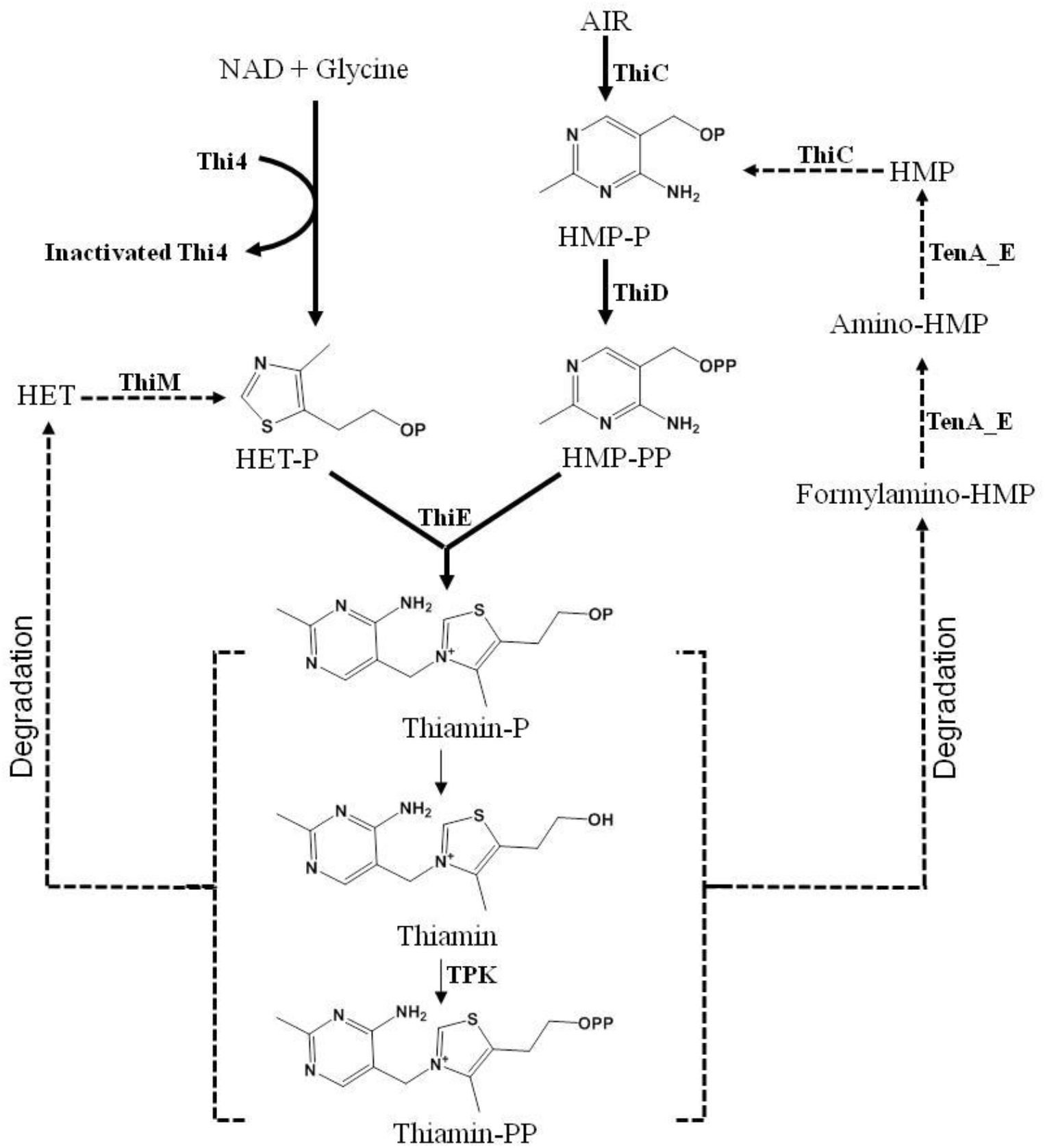
Thiamin *de novo* biosynthesis and salvage pathway in plants. ThiC, phosphomethylpyrimidine synthase; ThiD, phosphomethylpyrimidine kinase; ThiE, thiamin-phosphate pyrophosphorylase; Thi4, single turnover thiazole biosynthesis enzyme; TPK, thiamin pyrosphokinase. AIR, 5-aminoimidazole ribotide; HMP, 4-amino-5-hydroxymethyl-2-methylpyrimidine; Formylamino-HMP, N-formyl-4-amino-5-aminomethyl-2-methylpyrimidine; Amino-HMP, 4-amino-5-aminomethyl-2-methylpyrimidine; HMP-P, 4-amino-5-hydroxymethyl-2-methylpyrimidine phosphate; HMP-PP, 4-amino-5-hydroxymethyl-2-methylpyrimidine diphosphate; HET-P, 4-methyl-5-(β*-*hydroxyethyl)thiazole phosphate; ThiM, thiazole kinase; TenA_E, a thiaminase II protein family; Thiamin-P, thiamin monophosphate; Thiamin-PP, thiamin diphosphate.

Thiamin has been demonstrated to play a vital role in resistance against biotic and abiotic stress conditions in various plant species (Sayed and Gadallah, 2002; Ahn et al., 2005; Tunc-Ozdemir et al., 2009; Boubakri et al., 2013). Abiotic stresses such as drought, salinity, extreme temperatures, chemical toxicity and oxidative stress are serious threats to agriculture and are the major causes of crop loss worldwide. Furthermore, these stresses can cause morphological, physiological, biochemical and molecular changes which adversely affect plant growth and productivity (Wang et al., 2000). They can reduce the average yield of most staple crops by more than 50% (Boyer, 1982; Bray, 2000; Alcázar et al., 2006).

Several studies have shown that plants respond to abiotic stresses by increasing thiamin biosynthesis and accumulation. Tunc-Ozdemir et al. (2009) demonstrated that plants subjected to various abiotic stress treatments including high salt, drought, high light intensity, and paraquat synthesize elevated levels of thiamin. This stress-induced increase in thiamin levels has been shown to correlate with increased expression of thiamin biosynthetic genes. Additionally, Rapala-Kozik et al. (2008) showed the increased thiamin content of *Zea mays* seedlings under drought, high salt, and oxidative stress conditions was accompanied by an elevation in the activity of thiamin biosynthetic enzymes. Furthermore, it seems that this phenomenon is not limited to terrestrial plants. For instance, the accumulation of thiamin in various algal species under high light, salinity, and temperature stresses has been reported (Sylvander et al., 2013). These observations suggest an important role for thiamin in abiotic stress tolerance. In support of this idea, Sayed and Gadallah (2002) showed that the application of thiamin on the shoots and roots of salt-stressed sunflower plants alleviated the detrimental effects of salt stress on plant growth by improving both the cell membrane integrity and K^+^ uptake. Tunc-Ozdemir et al. (2009) later showed that the exogenous application of thiamin could protect *Arabidopsis* from paraquat-induced oxidative damage. The authors discuss the possibility that TPP could be a critical factor in plant stress tolerance, as its levels were higher than the free thiamin and TMP. This could be related to the role of TPP in the production of reductants, including NADH and NADPH, in order to combat oxidative stress damage (Tunc-Ozdemir et al., 2009; Rapala-Kozik et al., 2012). Furthermore, the exogenous application of thiamin can promote the antioxidant defense system of *Zea mays* at early stages of development (Kaya et al., 2015).

Transcriptome and proteome analysis were also performed in recent years to gain more insight about the relationship between thiamin biosynthesis and stress tolerance in plants. Transcriptomic studies have been reported on the accumulation of the transcripts of some enzymes in relation to thiamin biosynthesis under heat and drought stress conditions in plants (Rizhsky et al., 2004). Moreover, leaf proteome analysis of *Populus euphratica* plants subjected to heat stress showed changes in the abundance of thiamin biosynthesis enzymes (Ferreira et al., 2006). Proteomic analysis has also revealed that thiamin has an important role in cold stress response in rice seedlings (Cui et al., 2005).

Taken together, these evidences confirm that thiamin plays an important role to sustain normal functioning of plants subjected to various environmental stresses. Therefore, it is important to take advantage of biotechnology approaches to help plants overcome these environmental challenges in order to secure food supply worldwide.

The aim of this research is to generate metabolically engineered plants with improved thiamin content in the seeds in order to: 1-boost seed nutritional value, 2-increase abiotic stress tolerance and, 3-determine the possible impact of excess seed thiamin on various metabolic pathways in plants. To achieve these goals, we overexpressed *Arabidopsis Thi4*, *ThiC*, *ThiE*, and *TPK1* genes under the control of seed-specific promoters and analyzed the transgenic seeds of *Arabidopsis* and oilseed crop *C. sativa*. Ultimately, transgenic plants displayed improved nutritional content and increased abiotic stress tolerance, suggesting this strategy is a viable method for crop improvement. In addition, the transgenic seeds in both *Arabidopsis* and *C. sativa* produced more oil in comparison to the control seeds. Indeed, this is a novel finding which demonstrates that thiamin over-production is capable of shifting the carbon flux in plant seeds.

## RESULTS

### Overexpression of thiamin biosynthesis genes in wild type *Arabidopsis* and *C. sativa* seeds

We used *Brassica* napin and oleosin promoters to overexpress the *AtThi4*, *AtThiC* genes, respectively, and soybean glycinin promoter to overexpress *AtThiE* and *AtTPK1* genes in wild type *Arabidopsis* and *C. sativa* seeds (Fig. 2). These promoters were chosen because they are known to be expressed in both the endosperm and embryo tissues of the seeds from various dicot and monocot species (Keddie et al., 1994; Iida et al., 1995; Ellerström et al., 1996). More than twenty homozygous lines were obtained from both species. Eight independent homozygous lines of 3-gene transformant and seven independent homozygous lines of *AtTPK1* transformants were used for this study.

**Figure 2:**
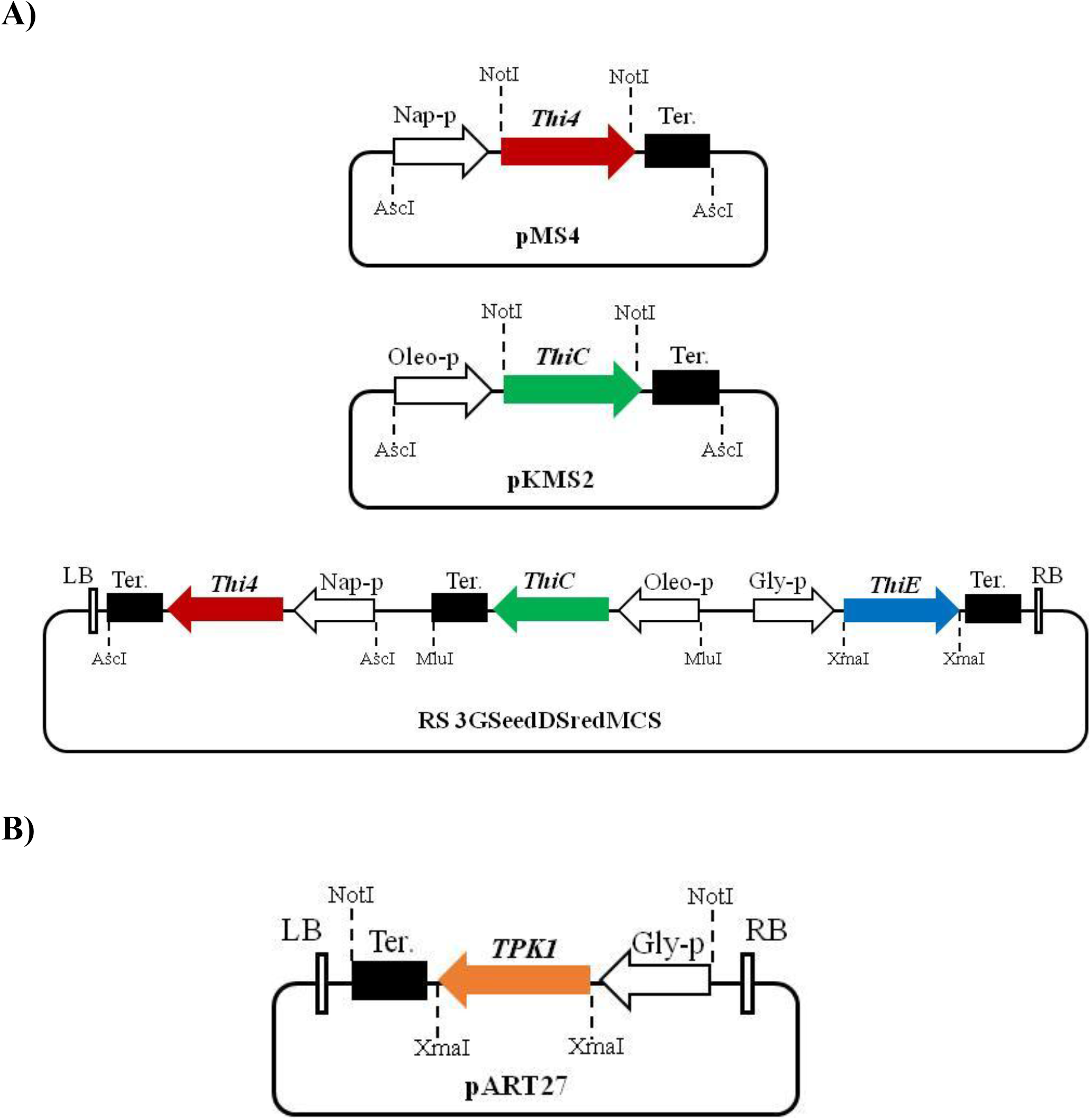
Vectors used for *Arabidopsis* and *C. sativa* transformation. **(A)** Simplified diagram depicting *Thi4* and *ThiC* gene cassettes in pMS4 and pKMS2 vectors. RS 3GSeedDSredMCS binary vector was used to clone *ThiE* gene. *Thi4* and *ThiC* gene cassettes were also sub-cloned into RS 3GSeedDSredMCS binary vector. MluI and AscI have compatible ends. (B) *TPK1* gene was first cloned into RS 3GSeedDSredMCS vector and the *TPK1* gene cassette was PCR amplified and cloned into pART27 transformation vector. Nap-p, Oleo-p, and Gly-p are napin, oleosin, and glycinin promoters, respectively. Ter.: terminator, LB: left border, RB: right border.

### Transgenic *Arabidopsis* and *C. sativa* plants accumulated greater thiamin in the seeds

Thiamin content analysis of control and transgenic lines by HPLC revealed that overexpression of *AtThi4*, *AtThiC*, and *AtThiE* genes under the control of seed-specific promoters significantly boosted the thiamin levels in the seeds. All transgenic lines had greater abundance of thiamin in comparison to the controls (Fig. 3A). The thiamin level in the *Arabidopsis* seeds ranged from 81 (line 6-5) to 131 ng per mg seed weight (line 13-15), which is approximately 4.2- to 7-fold greater than those in the controls. In addition, the thiamin content in the *C. sativa* seeds ranged from 27 (line 9-10) to 71 ng per mg seed weight (line 18-4), which is about 2.5- to 6.4-fold increase in comparison to the controls. Interestingly, thiamin in the form of cofactor (TPP) was not detectable in either transgenic or control seeds.

**Figure 3:**
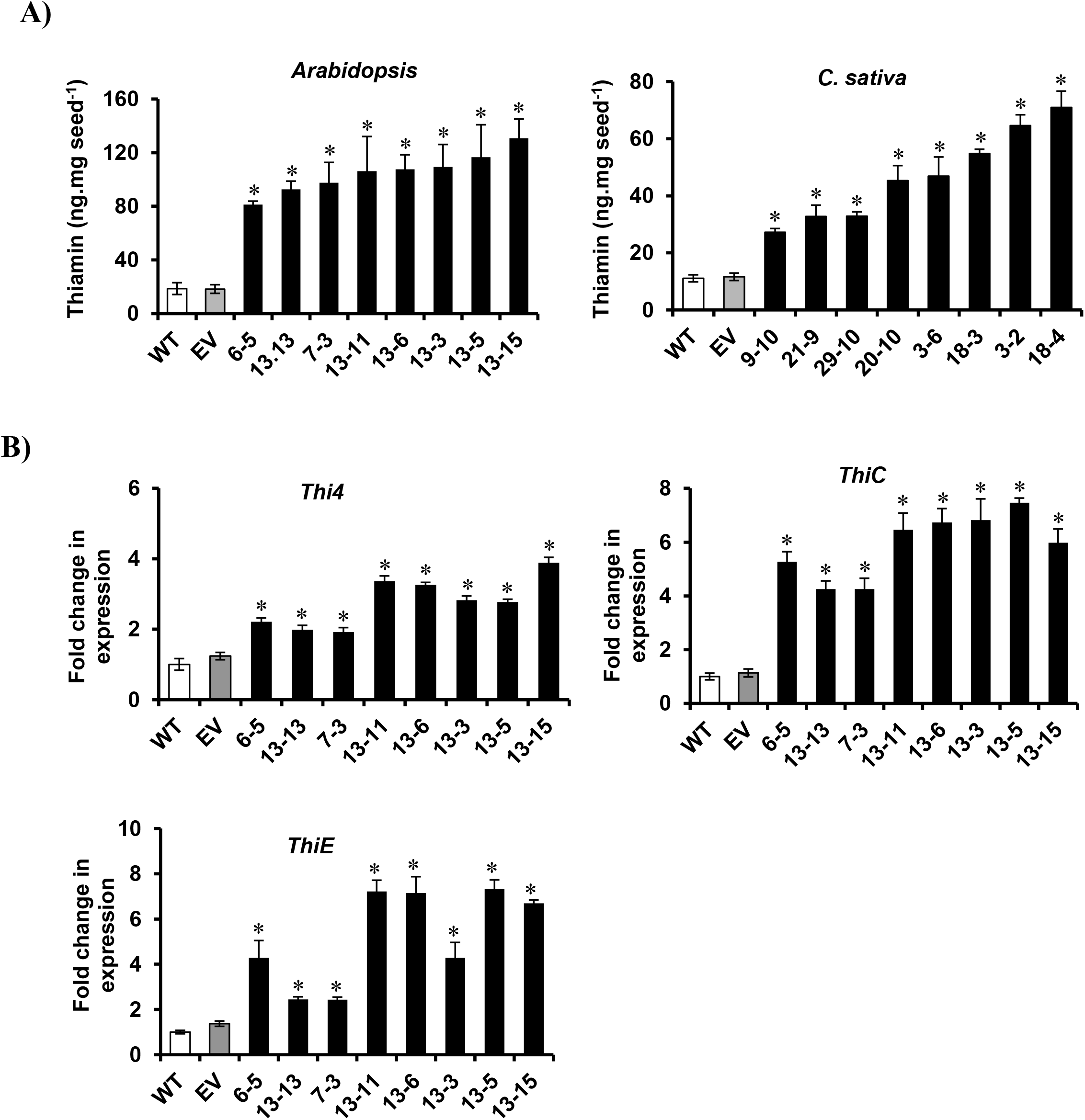
HPLC analysis showed thiamin over-accumulation in transgenic seeds. (A) Increased seed thiamin content of homozygous transgenic *Arabidopsis and C. sativa* plants (3-gene overexpressors). (B) Transcript abundance of thiamin biosynthetic genes in 3-gene overexpressing *Arabidopsis* plants. Values are mean ± SD. * indicates significant difference at *P*<0.05. WT: wild type, EV: empty vector control.

To evaluate the expression pattern of *Thi4*, *ThiC*, and *ThiE* genes in transgenic seeds, qRT-PCR was performed using 9-day-old developing siliques in *Arabidopsis*. The results showed that the transcripts of all three genes were significantly increased. The highest expression (>7-fold) was observed in the *ThiC* and *ThiE* genes, while the *Thi4* transcript abundance level was approximately 4-fold more than in the control plants (Fig. 3B).

### Transgenic *Arabidopsis* and *C. sativa* seedlings showed enhanced oxidative stress tolerance

To test the hypothesis that transgenic seedlings are more tolerant to oxidative stress, root growth assay was carried out using the seeds of transgenic and control lines grown on agar medium supplemented with various concentrations of paraquat. Root growth assay was chosen because it is fast, accurate, and easy to perform (Verslues et al., 2006; Ciftci-Yilmaz et al., 2007). Paraquat was used because it induces the production of superoxide radical, which has detrimental effects on the electron transfer mechanism in the chloroplast and mitochondrion (Bowler et al., 1991; Tunc-Ozdemir et al., 2009). The results showed that root growth rate (RGR) in transgenic *Arabidopsis* seedlings was significantly higher than in the control lines. The RGR for the transgenic seedlings grown on agar plates and supplemented with 0.05 and 0.1 µM paraquat showed an increase of approximately 1.3-fold compared to the control lines. For transgenic lines supplemented with 0.25 µM paraquat the increase was roughly 1.5-fold (Fig. 4A).

**Figure 4:**
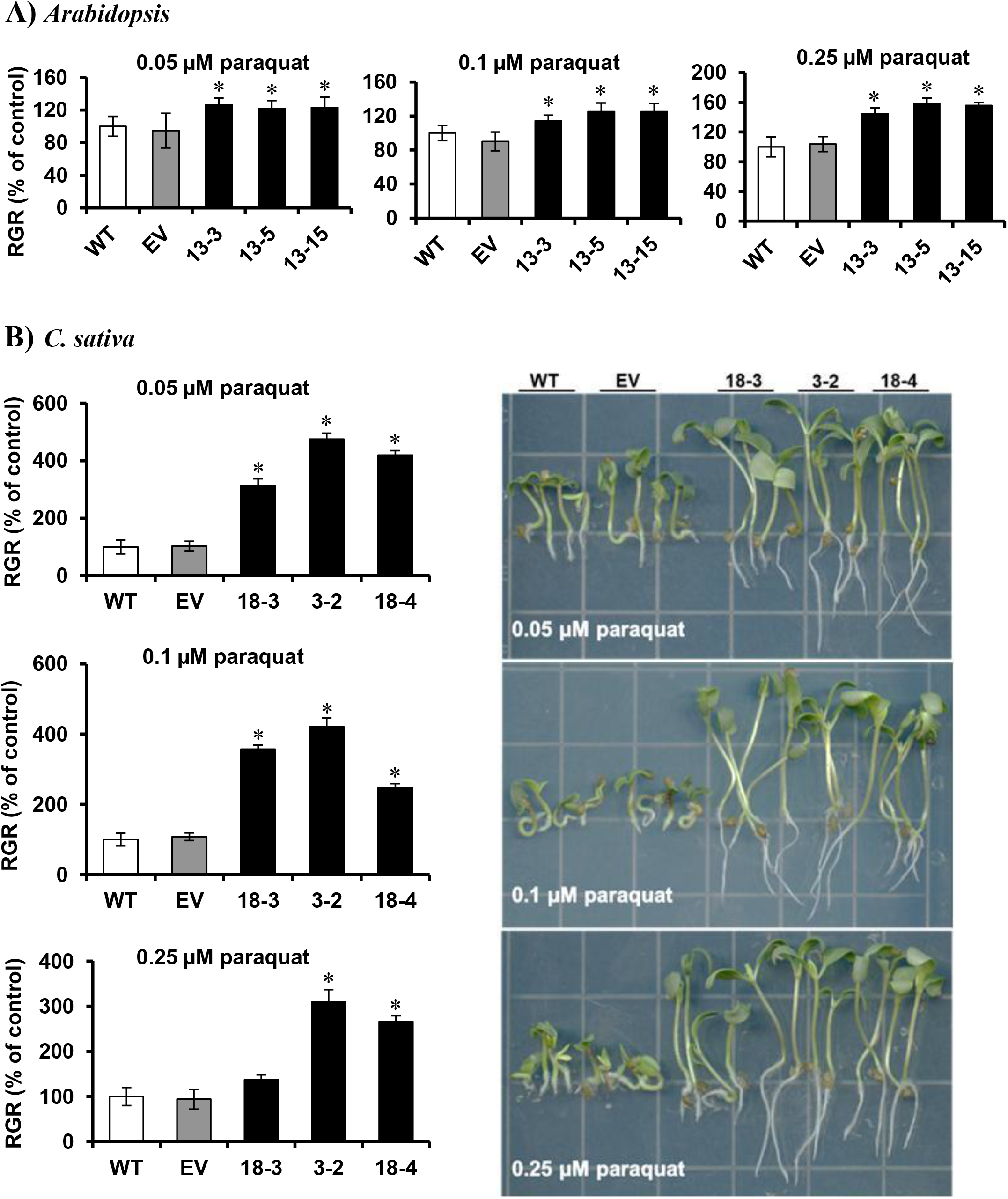
Transgenic lines are more resistant to oxidative stress. Effect of paraquat on root growth rate (RGR) of 9-day old *Arabidopsis* (A) and *C. sativa* (B) seedlings overexpressing 3 genes in the seed. In *Arabidopsis* 100% of the WT in 0.05, 0.1, and 0.25 µM paraquat equals 18.9, 14.2, and 6.6 mm, respectively. In *C. sativa* 100% of the WT in 0.05, 0.1, and 0.25 µM paraquat equals 0.5, 0.5, and 0.7 mm, respectively. Values are mean ± SE. * indicates significant difference at *P*<0.05. WT: wild type, EV: empty vector control.

On the other hand, the RGR in *C. sativa* transgenic seedlings (Fig. 4B) was much higher than the RGR in *Arabidopsis* transgenic seedling in all 3 different concentrations of paraquat. Compared to *C. sativa* control seedlings, the RGR for *C. sativa* transgenic seedlings in 0.05, 0.1, and 0.25 µM paraquat was 4.7-, 3.6-, and 3-fold higher, respectively. The line 18-3 showed about 1.3-fold increase in RGR, which was not statistically significant (Fig. 4B). These results suggest that increasing thiamin levels in seeds enhances resistance to paraquat induced oxidative stress of germinating seedlings.

### Elevation of seed thiamin levels improved germination rate and seedling viability under salt and oxidative stress conditions

To investigate how thiamin over-producing seeds react to the abiotic stress conditions during the onset of germination, the seeds of transgenic plants were grown on agar plates supplemented with various concentrations of NaCl and paraquat, separately. The results exhibited transgenic *Arabidopsis* and *C. sativa* seeds were more tolerant than the control seeds in the presence of 50 and 100 mM NaCl for *Arabidopsis* and 150 and 200 mM NaCl for *C. sativa* seeds (Fig. 5A and B). In *Arabidopsis*, under 50 mM salt in the medium more than 97% of the seeds in line 13-15 were germinated at day 2, compared with approximately 57% seed germination for the controls (Fig. 5A). The values for the seeds treated with greater salt stress (100 mM), were approximately 16 and 46% for the controls and transgenic line 13-15, respectively (Fig. 5A). While *C. sativa* transgenic seeds showed almost the same trend in germination rate in 150 and 200 mM salt, the control lines were highly sensitive to both concentrations especially to 200 mM salt, as the germination rate reached about 10% at day 2 in comparison to 55% in transgenic line 3-2. After day 2, the germination rate of transgenic seeds highly increased compared to the controls. For instance, at day 3, it reached 80-100% in both *Arabidopsis* and *C. sativa* transgenic lines while the germination in control lines remained lower, specifically in higher salt concentrations (Fig. 5A and B).

**Figure 5:**
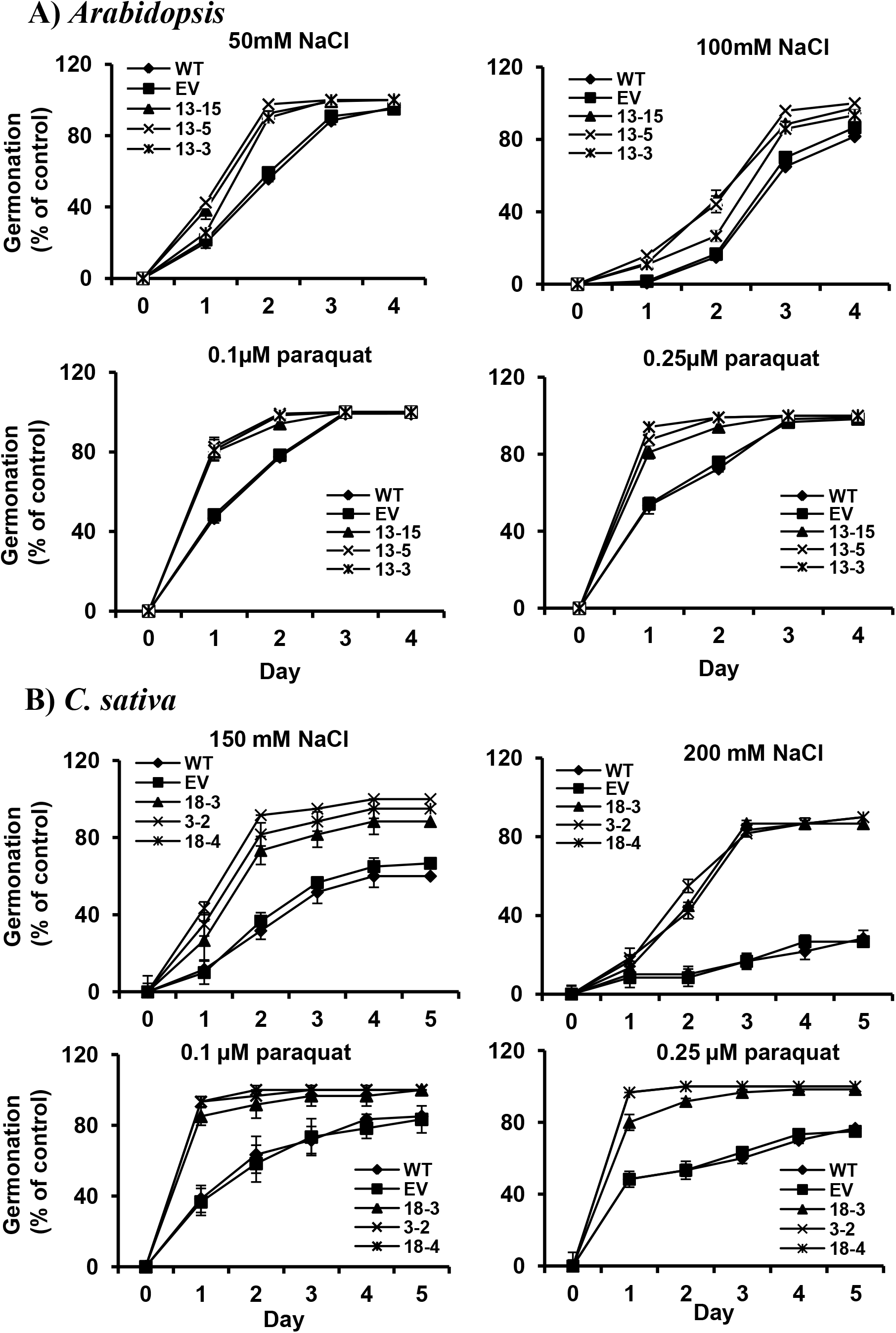
Transgenic seeds showed better germination under salt and paraquat stress. Effects of salt and paraquat on the germination of 3-gene overexpressing *Arabidopsis* (A) and *C. sativa* (B) seeds. Radicle emergence was used as a marker for seed germination. Germination assays were carried out using four replicates each consisting of 30-45 seeds. Bars denote mean ± SE. WT: wild type, EV: empty vector control.

A similar result was observed in the paraquat experiments. Germination of control seeds under paraquat treatment was inhibited significantly at day 1 for both *Arabidopsis* and *C. sativa*. The results showed that at day 1 about 46% of control seeds for *Arabidopsis* and roughly 37% for *C. sativa* were germinated compared to more than 80% germination in transgenic lines for both species grown on agar medium supplemented with 0.1 µM paraquat (Fig. 5A and B). In *Arabidopsis*, germination continued at day 2 with significant difference between the controls and transgenic seeds (approximately 78% in controls vs. 99% in line 13-5), then reached about 100% in all genotypes at day 3. Unlike *Arabidopsis*, the germination in *C. sativa* control seeds never reached 100%. The germination rate on the plates supplemented with 0.25 µM paraquat was approximately 50% for both *Arabidopsis* and *C. sativa* control seeds at day 1 while it reached to 94% and 97% for the high-thiamin lines in *Arabidopsis* and *C. sativa*, respectively (Fig. 5A and B). Seeds of all genotypes continued to germinate throughout day 2 to 5, although the germination rate of high-thiamin lines were significantly greater than those of the controls (Fig. 5A and B). Similar to 0.1 µM paraquat treatment, *C. sativa* control seeds never reached to 100% germination during the experiments (Fig. 5B).

To assess whether the seeds are able to survive and to generate seedlings after day 5 on agar plates supplemented with 100 mM salt and 0.25 µM paraquat, the corresponding plates were kept in the growth chamber for 9 days and the number of seedlings were counted. The results for *Arabidopsis* revealed that although about 88% of control seeds germinated by day 5 in both 100 mM salt and 0.25 µM paraquat treatments, only 18% of them were able to form seedlings under the 100 mM salt condition (Fig. 6A). In contrast, up to 47% of transgenic seeds generated seedlings. Under 0.25 µM paraquat treatment, 65 and 98% of control and high-thiamin lines were able to form seedling, respectively (Fig. 6A). In *C. sativa*, only 11% of the control seeds produced seedlings and survived in 150 mM NaCl; however, up to 35% of the seeds from the transgenic lines formed viable seedlings (Fig. 6B). In the plates containing 0.25 µM paraquat, 58% of the control seeds survived and generated seedlings compared to up to 89% in transgenic lines (Fig. 6B). Together, these results clearly indicate that thiamin accumulation mediated by the overexpression of *Thi4*, *ThiC*, and *ThiE* genes helps both *Arabidopsis* and *C. sativa* seeds to cope with abiotic stress.

**Figure 6:**
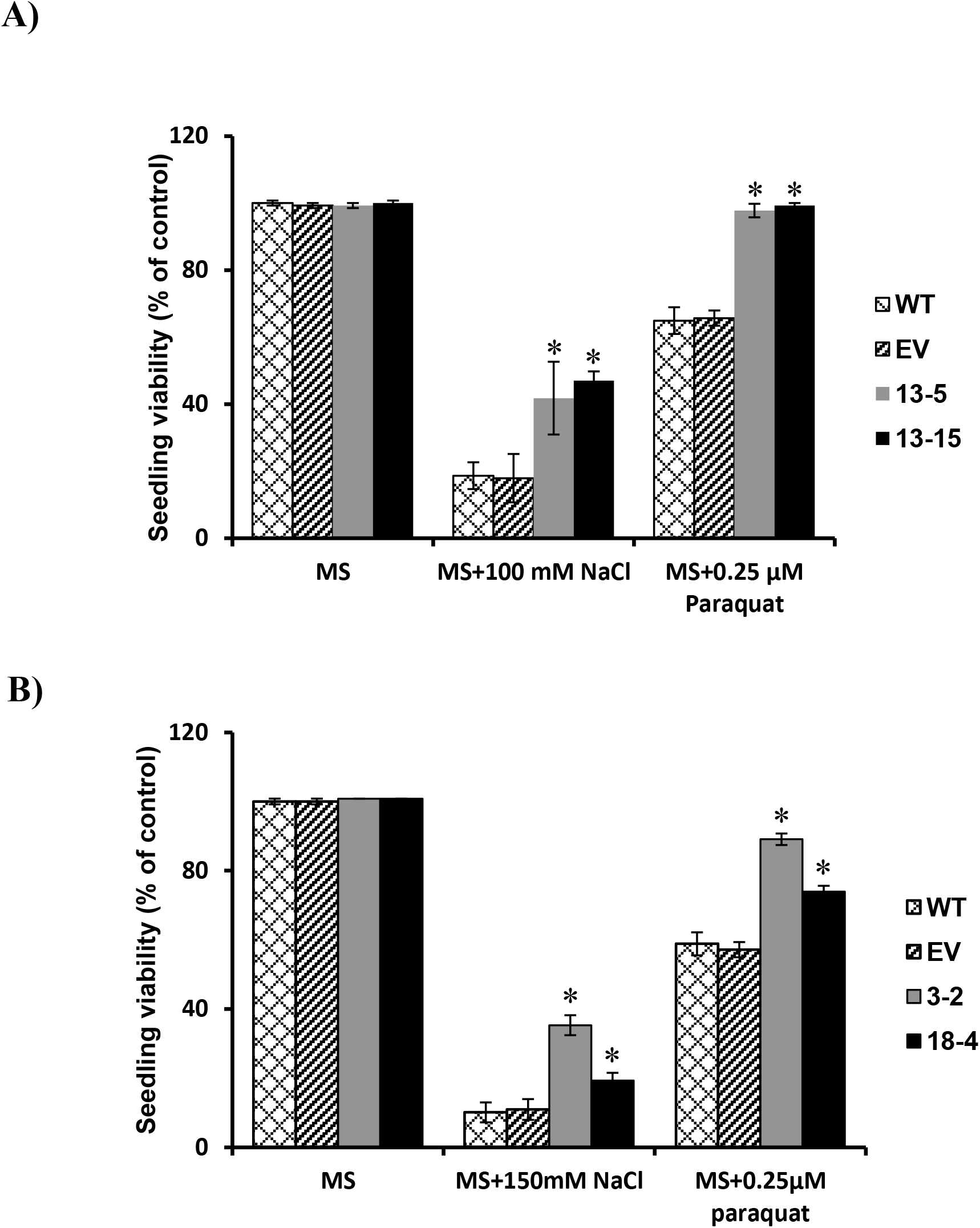
Greater seedling viability was observed in transgenic seeds under salt and paraquat treatments. Effects of salt and paraquat treatments on seedling viability of 3-gene overexpressing *Arabidopsis* (A) and *C. sativa* (B) seeds. The assays were performed in three replicates each consisting of 45 seeds. 100% of the wild type in NaCl and paraquat treatments were 8.3 and 29 seeds for *Arabidopsis*, and 4 and 23.3 for *C. sativa*, respectively. Bars denote mean ± SE. * indicates significant difference at *P*<0.05. WT: wild type, EV: empty vector control.

### Thiamin over-accumulation altered seed phenotype in transgenic plants

To assess whether over-production of thiamin alters the metabolic pathways in high-thiamin lines, seed oil, total protein, and carbohydrate content were measured. Seeds of *Arabidopsis* plants produce triacylglycerols (TAGs) which are the esterified forms of the fatty acids (Li et al., 2006). To evaluate the *Arabidopsis* seed oil content, we used the direct methylation method of intact seeds as described by Li et al. (2006), followed by gas chromatography (GC) analysis of fatty acid methyl esters (FAMEs). Surprisingly, we noticed that transgenic seeds accumulated more oil in the form of FAME than those of the control seeds. The levels of oil content per seed dry weight ranged from 42.1% in line 6-5 to 46.1% in line 13-15, which is 6.3-16.4% greater than the wild type seeds (Table 1). To the best of our knowledge, this is the first report regarding the effect of thiamin on fatty acid biosynthesis in plants. Additionally, the analysis of fatty acid composition in *Arabidopsis* seeds showed that palmitic acid (C_16_:0), stearic acid (C_18_:0), and Oleic acid (C_18_:1) levels were higher compared to the wild type seeds. However, the level of linolenic acid (C_18_:3) was lower than the wild type (Table 3).

**Table 1:** Seed compositions of transgenic *Arabidopsis* and *C. sativa* plants overexpressing *AtThiC, AtThi4*, and *AtThiE* genes (% per seed weight). Values are means of three biological replicates ± SE.

| *Arabidopsis* |  |  |  |  |  |  |
| --- | --- | --- | --- | --- | --- | --- |
| Genotypes | Lipid | *P*- value | Protein | *P*- value | Carbohydrate | *P*- value |
| WT | 39.6 ± 0.84 | | 35.2 ± 2.01 | | 7.2 ± 0.46 | |
| EV | 40.0 ± 0.56 | 0.88 | 36.3 ± 1.40 | 0.98 | 7.3 ± 0.52 | 0.65 |
| **6-5** | **42.1 ± 0.61** | 0.04 | **14.8 ± 0.69** | 0.0001 | **9.8 ± 0.42** | 0.001 |
| **13-13** | **43.7 ± 0.56** | 0.004 | **8.7 ± 0.96** | 0.0004 | **9.9 ± 0.74** | 0.03 |
| **7-3** | **43.4 ± 0.03** | 0.0001 | **14.4 ± 0.53** | 0.0008 | **10.5 ± 0.43** | 0.0008 |
| **13-11** | **43.1 ± 1.00** | 0.002 | **13.7 ± 0.81** | 0.0003 | **10.7 ± 0.22** | 0.0002 |
| **13-6** | **43.9 ± 0.78** | 0.01 | **18.1 ± 0.50** | 0.0004 | **11.3 ± 0.76** | 0.01 |
| **13-3** | **44.8 ± 0.60** | 0.002 | **12.3 ± 1.95** | 0.001 | **11.7 ± 0.35** | 0.0005 |
| **13-5** | **43.9 ± 0.70** | 0.01 | **13.0 ± 0.64** | 0.0006 | **11.4 ± 0.11** | 0.0002 |
| **13-15** | **46.1 ± 0.71** | 0.003 | **13.3 ± 0.10** | 0.0002 | **11.6 ± 0.93** | 0.02 |
| *C. sativa* |  |  |  |  |  |  |
| WT | 31.8 ± 0.11 | | 26.2 ± 1.00 | | 5.9 ± 1.86 | |
| EV | 32.1 ± 0.19 | 0.16 | 28.2 ± 0.90 | 0.11 | 6.0 ± 1.69 | 0.49 |
| **9-10** | **36.3 ± 0.47** | 0.0004 | **19.3 ± 1.19** | 0.006 | **16.5 ± 1.51** | 0.01 |
| **21-9** | **34.6 ± 0.13** | 0.001 | **17.8 ± 1.10** | 0.002 | **15.0 ± 2.02** | 0.02 |
| **29-10** | **33.6 ± 0.33** | 0.004 | **20.3 ± 1.60** | 0.02 | **11.0 ± 0.41** | 0.03 |
| **20-10** | **34.3 ± 0.59** | 0.01 | **16.9 ± 0.80** | 0.001 | **19.1 ± 1.58** | 0.003 |
| **3-6** | **37.5 ± 0.12** | 0.0002 | **16.7 ± 1.93** | 0.01 | **14.6 ± 3.42** | 0.04 |
| **18-3** | **35.0 ± 0.38** | 0.001 | **16.1 ± 2.17** | 0.01 | **15.1 ± 1.71** | 0.01 |
| **3-2** | **36.7 ± 0.36** | 0.0001 | **19.6 ± 0.63** | 0.003 | **13.9 ± 0.84** | 0.01 |
| **18-4** | **37.2 ± 0.58** | 0.0004 | **15.1 ± 0.55** | 0.0003 | **19.2 ± 0.79** | 0.001 |

WT: wild type, EV: empty vector control.

A striking difference in the amount of total protein was observed between high-thiamin lines and controls (Table 1). For instance, the high-thiamin line 13-15 showed about 22% decrease in total protein content per seed dry weight compared to the wild type seeds. In contrast, the total carbohydrate levels were elevated. The level of total sugar in line 13-15 was 4.4% greater per seed dry weight than the wild type controls (Table 1). Moreover, soluble sugars and seed weight analyses exhibited that thiamin over-producing seeds contained significantly more soluble sugars and had greater weight than the control seeds (Fig.7).

**Figure 7:**
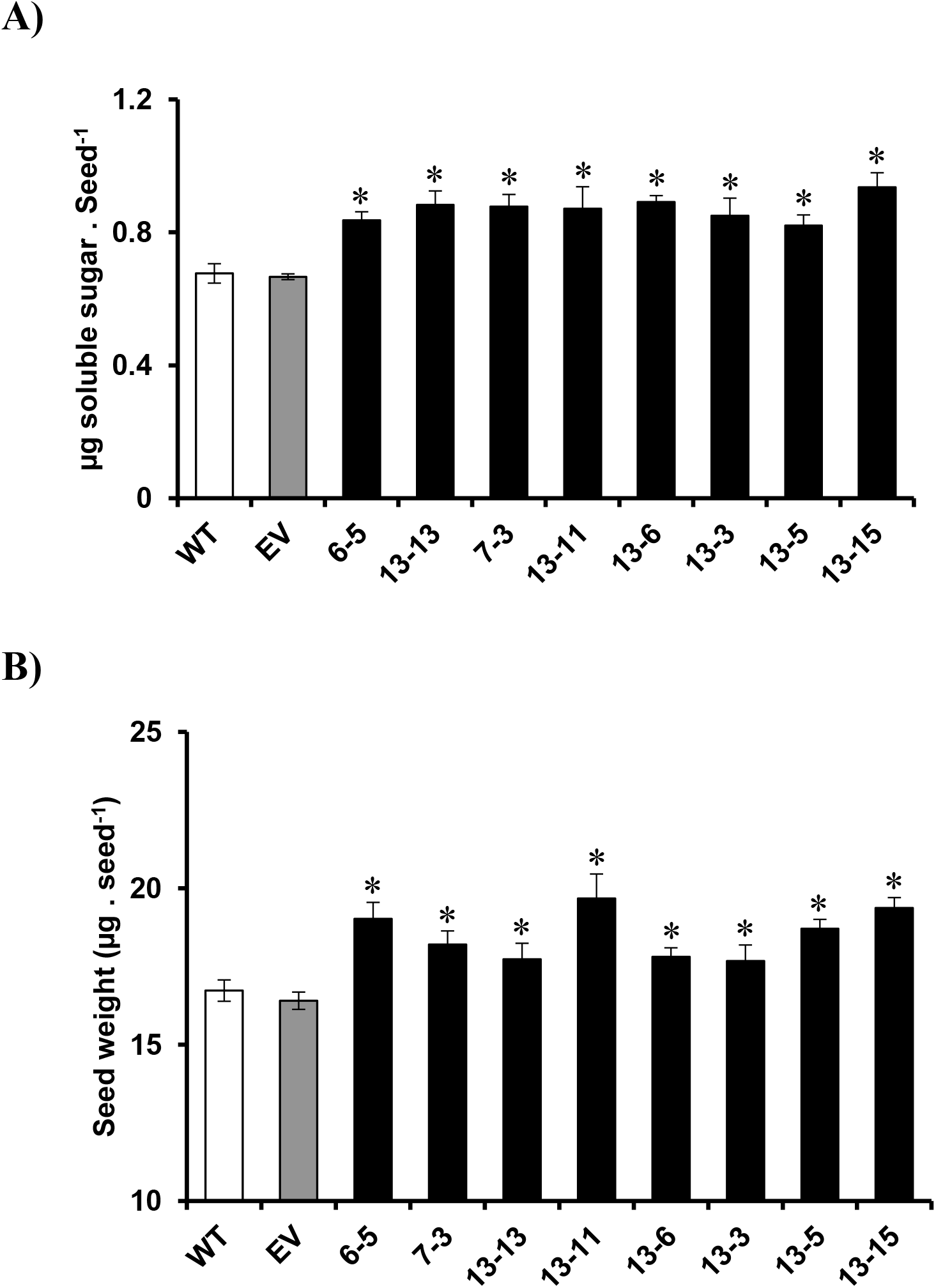
Overexpression of thiamin biosynthetic genes resulted in an increase in soluble sugars content and seeds mass. (A) Seed soluble sugars content of controls and 3-gene overexpressing *Arabidopsis*. (B) Seed weight of controls and 3-gene overexpressing *Arabidopsis* using 100 seeds for each line in triplicates. Values are mean ± SE. * indicates significant difference at *P*<0.05. WT: wild type, EV: empty vector control.

Interestingly, the *C. sativa* seed oil content in transgenic lines was also more than in wild type control seeds. The levels of oil per seed dry weight ranged from 33.6% in line 2-10 to 37.5% in line 3-6, which is 5.7-17.9% more than the wild type seeds (Table 1). *C. sativa* protein and total sugar levels showed the same trends as observed in *Arabidopsis* seeds.

Altogether, these results suggest that thiamin over-accumulation is able to shift the cell carbon flux between major metabolic pathways in *Arabidopsis* and *C. sativa* seeds.

### Thiamin over-production dramatically altered the gene expression levels of TPP-dependent enzymes in *Arabidopsis* seeds

To address whether over-production of thiamin in transgenic seeds is capable of changing the gene expression pattern of TPP-requiring enzymes, qRT-PCR was carried out for TKs, PDHs, α-KGDHs, and 1-deoxy-D-xylulose-5-phosphate synthases (DXPSs) using 9-day-old developing *Arabidopsis* siliques. The results indicated that transcript abundance levels in high-thiamin lines for all aforementioned enzymes were significantly increased by up to 2.5-fold compared to the wild type controls (Fig. 8).

**Figure 8:**
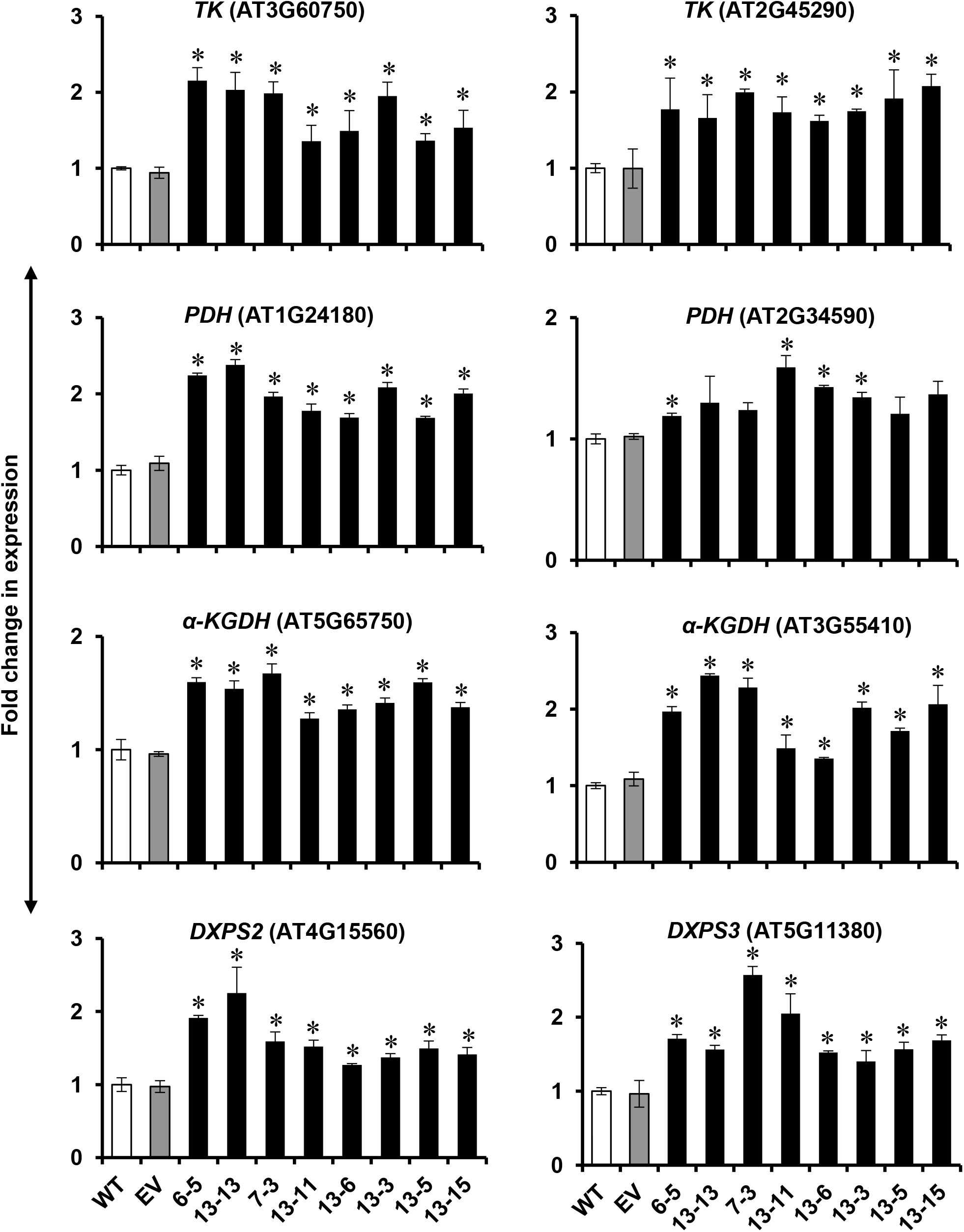
Transcript levels of TPP-dependent enzymes up-regulated by thiamin over-production in *Arabidopsis* seeds. *TK*: transketolase, *PDH*: pyruvate dehydrogenase, *α-KGDH*: α-ketoglutarate dehydrogenase, *DXPS*: deoxy-xylulose phosphate synthase. Bars represent means of three biological replicates ± SD. * indicates significant difference at *P*<0.05. WT: wild type, EV: empty vector control.

These results suggest that the accumulation of thiamin in transgenic seeds plays a significant role in up-regulation of TPP-dependent enzymes which are crucial for the generation of reducing molecules required for various metabolic pathways.

### Seed reserves in thiamin over-producing lines enhanced hypocotyl elongation

The necessity of thiamin for plants growth and development raised the question of whether the high-thiamin phenotypes in transgenic seeds would help their germination and growth on a medium which is lacking in thiamin and sucrose. To address this question, hypocotyl elongation assay was performed. The results revealed that the hypocotyl elongation in high-thiamin lines were affected by their higher thiamin pools when grown on MS plates without thiamin and sucrose, as they could form longer hypocotyls compared to the controls (Fig. 9). As shown in Fig. 9A, although the hypocotyl elongation continued from day 2 through day 6 for both transgenic and control *Arabidopsis* seeds, the transgenic lines formed longer hypocotyls than the controls. In line 13-15 for instance, at day 2 hypocotyl length was approximately 62% longer than the controls. This value for day 4, 6, and 8 was 43, 43, and 30%, respectively. After day 6, hypocotyl elongation reached to plateau for all genotypes (Fig. 9A). In *C. sativa*, significant differences in hypocotyl length between controls and high-thiamin lines were observable from day 3 of the experiment, and for the rest of the days were almost the same as we noticed in *Arabidopsis* (Fig 9 B). These results suggest that high-thiamin pool in transgenic seeds helps them to take advantage of their higher oil and sugar reserves to grow better when photosynthesis is blocked in darkness.

**Figure 9:**
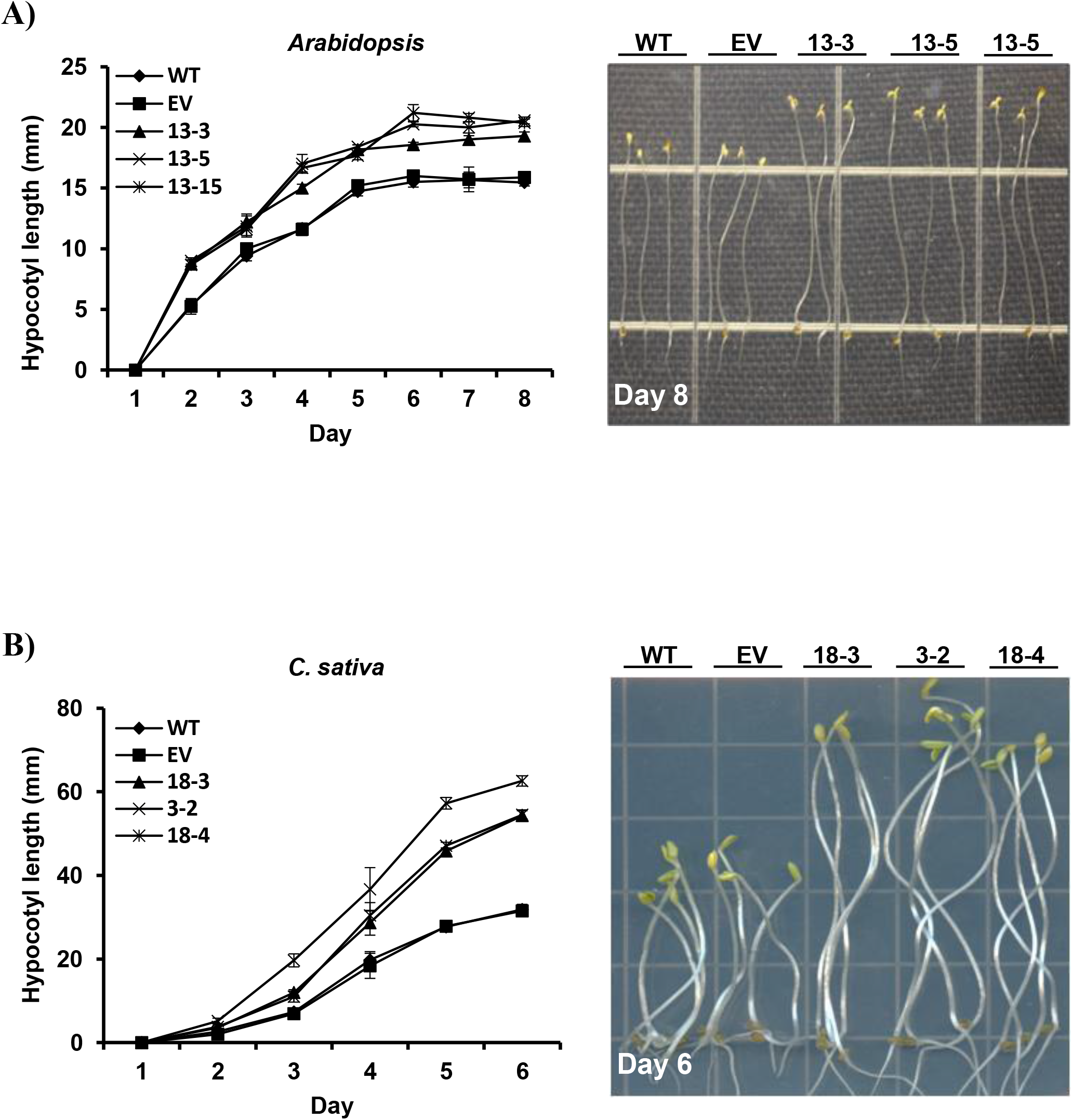
Hypocotyl elongation boosted in thiamin over-producing lines. Hypocotyl elongation was measured in *Arabidopsis* (A) and *C. sativa* (B) seeds. Seeds were plated on agar plates containing half strength MS salt without thiamin and sucrose. The plates were kept in 4 °C for 2 days in complete darkness. Plates were then placed in 22 °C under 100 µmol m^-2^ S^-1^ constant light for 24 h. Then, the plates wrapped in aluminum foil individually and placed vertically in 22 °C. The hypocotyl length was measured for 8 days for *Arabidopsis* and 6 days for *C. sativa*.

### *Arabidopsis* seeds overexpressing *TPK1* accumulated both free thiamin and TPP

In order to evaluate if TPP is responsible for the seed phenotypes results obtained from the overexpression of 3 genes (*Thi4*, *ThiC*, and *ThiE*), the *TPK1*, which is the last gene in thiamin biosynthesis pathway (Fig. 1) for the conversion of free thiamin to TPP was overexpressed under the control of the glycinin promoter (Fig. 2). We chose the *TPK1* because of its stronger expression in various parts of the *Arabidopsis* than *TPK2* (Ajjawi et al., 2007a). The homozygous transgenic and empty vector lines were generated as described for the 3-gene transformants, and the 7 best lines were used for further analysis. HPLC results showed that unlike the control seeds, *TPK1* transgenic lines could produce detectable amounts of TPP (Fig. 10A). The TPP content in transgenic seeds ranged from 19 to 20 ng per mg seed weight in lines 16-1 and 13-3, respectively. The total thiamin content in *TPK1* transgenic lines was about 2-fold greater than the wild type control (Fig. 10A). Additionally, qRT-PCR analysis showed that *TPK1* transcript abundance in transgenic lines was approximately 4- to 7-fold greater than the controls (Fig. 10B).

**Figure 10:**
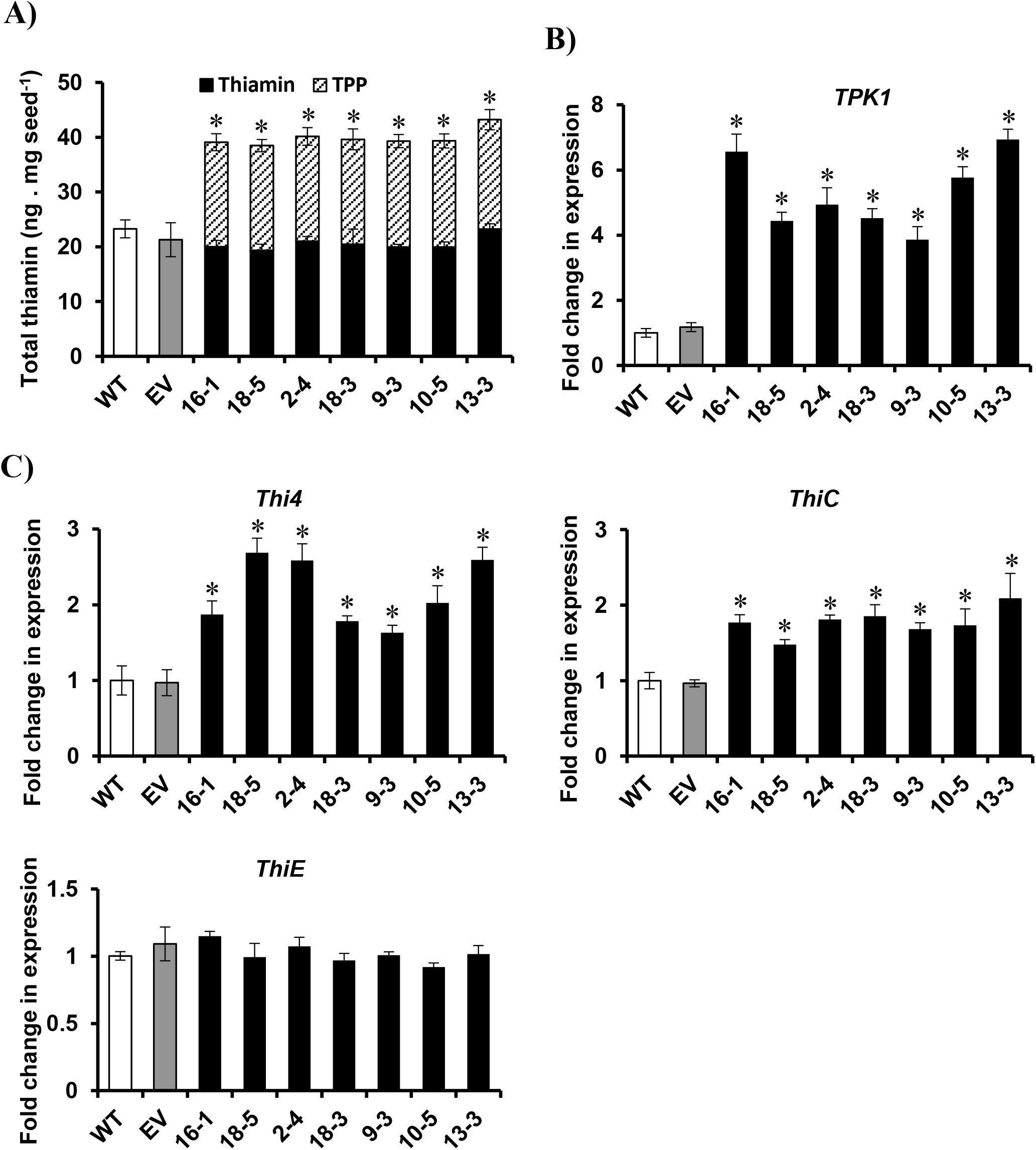
Overexpression of *TPK1* gene in *Arabidopsis* seeds resulted in the accumulation of both free thiamin and TPP. (A) HPLC analysis of seed thiamin content of homozygous transgenic *Arabidopsis* plants overexpressing *TPK1* gene. (B) Transcript abundance in *TPK1* overexpressing *Arabidopsis* plants. (C) Transcript abundance of thiamin biosynthetic genes in *TPK1* overexpressing *Arabidopsis* seeds. Values are means of three biological replicates ± SD. * indicates significant difference at *P*<0.05. WT: wild type, EV: empty vector control.

### TPP-overproduction in *Arabidopsis* seeds up-regulated the upstream genes involved in thiamin biosynthesis pathways

As shown in Fig. 10A, the total amounts of free thiamin and TPP in *TPK1* transgenic lines was about 2-fold greater than in the controls. These results raised the question of whether the overexpression of *TPK1* gene is capable of altering the expression levels of the other genes involved in thiamin biosynthesis pathways. To address this question, qRT-PCR was carried out to determine the *Thi4*, *ThiC*, and *ThiE* genes expression levels in transgenic and control seeds. The results indicated that the expression of *Thi4* and *ThiC* genes affected by the overexpression of *TPK1* gene, as they were up-regulated up to 2.5-times more than that of the controls. On the contrary, no significant changes were observed in *ThiE* gene expression level (Fig. 10C). These results suggest that the overexpression of *TPK1* in *Arabidopsis* seeds transcriptionally regulates the upstream genes to provide enough free thiamin for TPP biosynthesis.

### *Arabidopsis* seeds overexpressing the *TPK1* exhibited altered storage products

To confirm the results regarding seed phenotypes observed in the 3-gene overexpressing lines and to find whether thiamin and/or TPP were responsible for those results, we analyzed the seed composition of *TPK1* transgenic seeds. Similarly, oil and carbohydrate levels were boosted in *TPK1* transgenic seeds. The GC results showed that although it was in lesser amount than 3-gene transformants, the oil level of *TPK1* transgenic seeds were elevated up to 10% (line 13-3) compared to control lines (Table 2). In addition, fatty acid composition of these seeds showed that they contain more Oleic acid (C_18_:1) than wild type seeds (Table 3). The sugar level was also increased up to 50% (line 2-4) than the controls. Consistent with the results for protein level in 3-gene overexpressing lines, *TPK1* transgenic plants stored roughly 60% less protein in their seeds (lines 18-3) than the wild type control (Table 2). These results clearly show that TPP can significantly influence carbon assimilation and partitioning between seed storage compounds in *Arabidopsis* seeds.

**Table 2:** Seed compositions of transgenic *Arabidopsis* plants overexpressing *TPK1* gene (% per seed weight). Values are means of three biological replicates ± SE.

| **Genotypes** | **Lipid** | ***P*- value** | **Protein** | ***P*- value** | **Carbohydrate** | ***P*- value** |
| --- | --- | --- | --- | --- | --- | --- |
| WT | 40.3 ± 0.31 | | 41.2 ± 3.61 | | 12.5 ± 1.45 | |
| EV | 40.8 ± 0.36 | 0.16 | 44.9 ± 5.22 | 0.33 | 12.8 ± 0.27 | 0.27 |
| **16-1** | **42.8 ± 0.75** | 0.02 | **28.4 ± 4.84** | 0.004 | **16.8 ± 0.10** | 0.01 |
| **18-5** | **41.4 ± 0.65** | 0.11 | **33.4 ± 2.13** | 0.01 | **17.9 ± 0.10** | 0.002 |
| **2-4** | **43.1 ± 0.58** | 0.006 | **25.8 ± 3.49** | 0.0001 | **18.8 ± 0.20** | 0.04 |
| **18-3** | **42.2 ± 0.30** | 0.006 | **25.1 ± 1.67** | 0.0009 | **14.3 ± 0.21** | 0.05 |
| **9-3** | **43.4 ± 0.92** | 0.02 | **27.8 ± 2.40** | 0.002 | **15.9 ± 0.52** | 0.007 |
| **10-5** | **42.8 ± 0.14** | 0.001 | **31.1 ± 2.96** | 0.01 | **16.4 ± 0.61** | 0.004 |
| **13-3** | **44.4 ± 1.10** | 0.01 | **31.3 ± 3.10** | 0.02 | **16.1 ± 0.55** | 0.006 |

WT: wild type, EV: empty vector control.

**Table 3:**
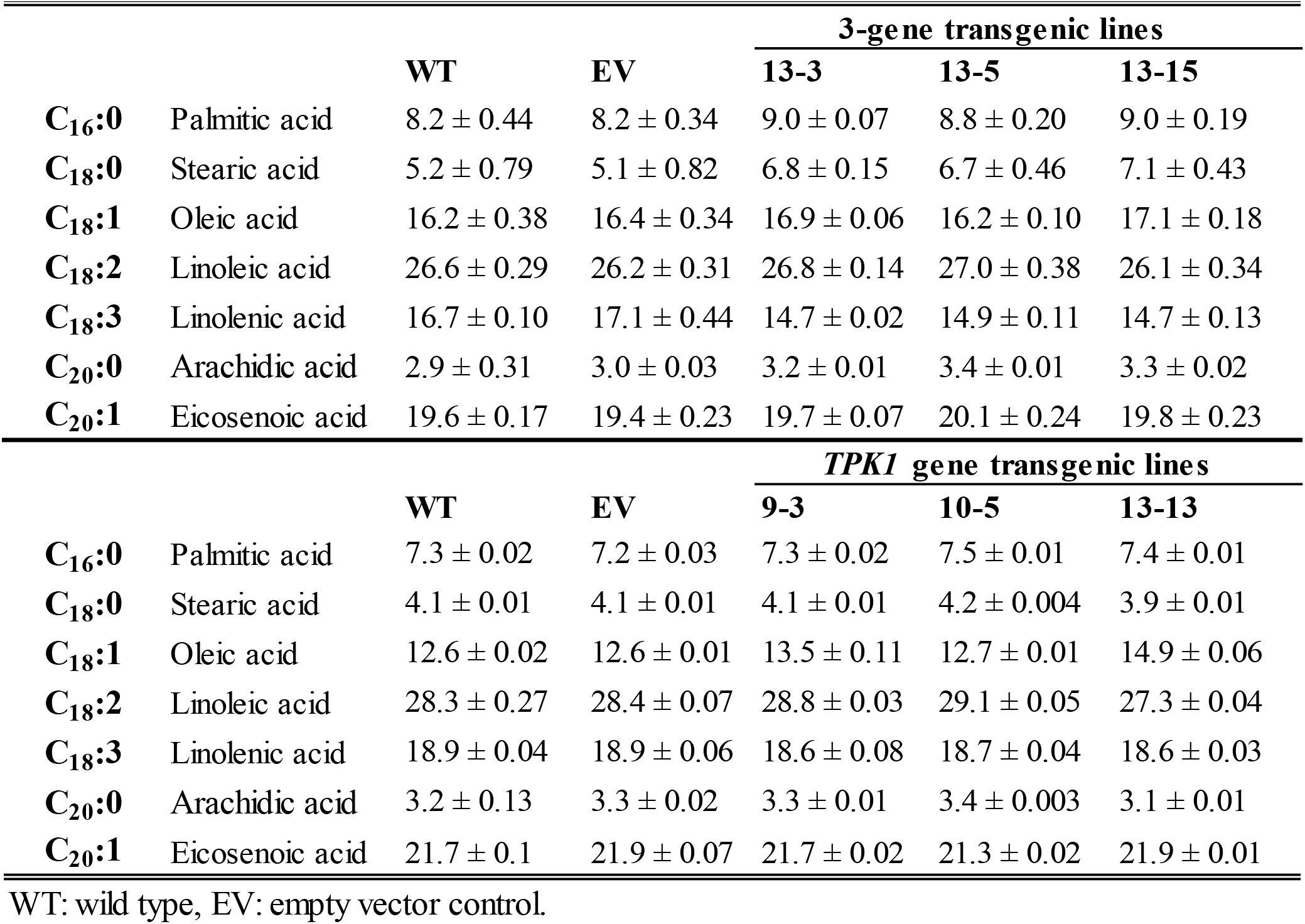
Fatty acid composition of the seeds (% of total fatty acid) from individual plants of each genotype. Values are means of three biological replicates ± SE.

## DISCUSSION

Thiamin cofactor, TPP, plays a key role in the metabolism of living organisms through the participation in metabolic pathways including Calvin cycle, acetyl-CoA formation, TCA cycle, pentose phosphate pathway (PPP), and the branched-chain amino acid biosynthesis (Friedrich, 1987). Although thiamin biosynthesis, salvage pathways, and regulation mechanisms have recently been well-characterized in bacteria and plants (Begley et al., 1999; Nosaka, 2006; Wachter et al., 2007; Ajjawi et al., 2007a; Yazdani et al., 2013; Zallot et al., 2014), there have been fewer reports regarding the generation of transgenic plants with elevated levels of thiamin. In this study, we showed that by using strong seed-specific promoters to co-overexpress the *AtThi4*, *AtThiC*, and *AtThiE* genes, the free thiamin level in the seeds of both *Arabidopsis* and *C. sativa* was boosted significantly (Fig. 3A). This suggests that overexpressing these three genes is sufficient to elevate the thiamin level in plant seeds. On the other hand, overexpression of the *TPK1* also led to the accumulation of TPP in transgenic seeds (Fig. 10A). Indeed, *TPK1* overexpression could up-regulate *Thi4* and *ThiC* genes (Fig. 10C) in order to increase free thiamin to the level required for synthesizing more TPP in the seed. These findings indicate that *Thi4* and *ThiC* genes are limiting in the thiamin biosynthesis pathway (Pourcel et al., 2013).

Our data show that both control and 3-gene overexpressing lines prefer to store thiamin in the form of free thiamin in the seeds (Fig. 3A). These results are in agreement with the previously reported evidence indicating the absence of phosphorylated forms of thiamin in mature seeds of some monocots and dicots such as rice, corn, soybean, and pea (Yusa, 1961; Molin et al., 1980; Gołda et al., 2004).

Oxidative stress is the major cause of damage for plants under environmental stress conditions because of the imbalance between reactive oxygen species (ROS) production and the activity of antioxidant systems (John and Scandalios, 1993). When plants are exposed to abiotic stress conditions, they switch on some genes that increase the level of certain metabolites which combat these stresses. Thiamin has been proven to be one of the metabolites that enhances tolerance to abiotic stresses such as heat, cold, salinity, and paraquat in plants (Tunc-Ozdemir et al., 2009). Although several studies in animal cells (Lukienko et al., 2000; Bâ, 2008) and yeast (Wolak et al., 2014) have given credit to thiamin as a potent antioxidant, there is not enough evidence to support this idea in plants. Most of the data regarding the role of thiamin in biotic and abiotic stress resistance relies on the addition of exogenous thiamin to the medium (Conrath et al., 2002; Sayed and Gadallah, 2002; Ahn et al., 2005; Tunc-Ozdemir et al., 2009), suggesting an indirect role for thiamin in scavenging ROS in plants by providing NADH and NADPH to combat oxidative stress (Tunc-Ozdemir et al., 2009; Asensi-Fabado and Munné-Bosch, 2010). In our study, the root growth assay showed that genetically engineered *Arabidopsis* and *C. sativa* seeds (3-gene overexpressing lines) had significantly increased RGR than that of the controls under paraquat treatment (Fig. 4). Paraquat is a well-known herbicide used to control weeds in agriculture (Babbs et al., 1989). As a powerful ROS producer, it can cause loss of photosynthetic activity via inhibition of ferredoxin reduction, NADPH generation, and regeneration of antioxidant enzymes (Lascano et al., 2012), as well as loss of cell membrane integrity (Kunert and Dodge, 1989). The increased RGR in transgenic *Arabidopsis* and *C. sativa* seedlings might be due to the elevation of free thiamin pool which can be subsequently converted to TPP cofactor required for the production of NADH and NADPH to reduce ROS levels in the cells (Tunc-Ozdemir et al., 2009).

Plants under stress conditions have a high demand for TPP, which activates enzymes such as TK, PDH, and α-KGDH to produce the additional NADH and NADPH required for scavenging the paraquat-generated ROS (Tunc-Ozdemir et al., 2009; Rapala-Kozik et al., 2012). In our study, it is also noteworthy that in thiamin over-producing lines, the *TK*, *PDH*, and *α-KGDH* gene expression were up-regulated (Fig. 8). This is consistent with the fact that TPP-dependent enzymes have a high demand for thiamin cofactor to produce reducing molecules in stress conditions (Rapala-Kozik et al., 2008). These reducing molecules are vital for the activity of several NAD(P)H-requiring enzymes such as glutathione reductase and mono-dehydro-ascorbate reductases, as well as the recycling of the oxidized form of vitamin E, which plays a crucial role in combating oxidative stresses (Arora et al., 2002; Miller et al., 2010). Furthermore, Tunc-Ozdemir et al. (2009) reported that the application of thiamin alleviated paraquat sensitivity in both wild type and *ascorbate peroxidase1* (*apx1*) mutants, which are highly sensitive to oxidative damage. These results suggest that thiamin can complement the function of APX1 enzyme and that thiamin may have a direct role in reducing oxidative stresses (Tunc-Ozdemir et al., 2009).

Interestingly, our transgenic seeds showed the greater total and soluble sugar content (Table 1 and Fig. 7A, respectively) compared to the control seeds, possibly due to the higher activity of TK enzyme in Calvin cycle, as indicated by a significant increase in *TK* gene expression level in transgenic seeds (Fig. 8). These results suggest an important role for increased carbohydrate levels in transgenic seeds in abiotic stress conditions. Hare et al. (1998) reported that accumulation of sugar is a common response of organisms to abiotic stress and can function as osmoprotectant and biomolecule stabilizer. Additionally, Kerepesi and Galiba (2000) observed that wheat plants accumulate more water-soluble carbohydrates during osmotic and salt stresses. Although the role of some soluble sugars, including glucose and fructose, in stress tolerance is controversial (Kerepesi and Galiba, 2000), it has been shown that other soluble carbohydrates, such as sucrose and fructans, have a potential role in adaptation to drought and salt stresses (McKersie and Lesheim, 2013). Sucrose has been shown prevent structural changes in soluble proteins in abiotic stress conditions (Kerepesi and Galiba, 2000). However, the exact mechanism of fructans in conferring stress resistance is not clear (Pilon-Smits et al., 1995). In addition, our results showed a significant up-regulation of the *DXPS* gene in thiamin over-producing lines (Fig. 8). DXPS enzyme can also serve as a defense agent in high-thiamin lines against ROS through the production of carotenoids in chloroplasts. Carotenoids are a vital class of antioxidants and play an important role in detoxifying the ROS within thylakoid membranes (Asensi-Fabado and Munné-Bosch, 2010).

Seed germination was also affected by both salt and paraquat treatments, and the transgenic lines showed greater rate of germination than the controls (Fig. 5). In addition, high-thiamin lines exhibited increased seedling viability under salt and paraquat treatments (Fig. 6). Inhibition of seed germination could be attributed to osmotic stress and ion toxicity caused by NaCl (Huang and Redmann, 1995). Likewise, Agarwal and Pandey, (2004) reported that salinity has detrimental effects on seed germination and seedling root and shoot length. Osmotic stress caused by salt stress reduces stromal volume in chloroplasts as well as generates ROS, which both play a critical role in photosynthesis inhibition (Price and Hendry, 1991). Oxidative damage generated by ROS can be alleviated by antioxidant systems which elevate plant tolerance to salt stress. Yoshimura et al. (2004) reported that overexpression of *Chlamydomonas* glutathione peroxidase in the cytosol and chloroplast of the tobacco plants boosted tolerance to oxidative stress induced by salt and paraquat treatments. This evidence would suggest that the excess amounts of thiamin in our transgenic lines could help regeneration of antioxidant systems by providing more reducing agents to alleviate the oxidative stress caused by salt and paraquat.

We used *Arabidopsis* and *C. sativa* in this study because *Arabidopsis* is a model plant for biology research (Meyerowitz and Somerville, 1994; Li et al., 2006) and both *Arabidopsis* and *C. sativa* are also close relative of rapeseed (*Brassica napus*), which is an important oilseed crop (Fröhlich and Rice, 2005; Li et al., 2006). We observed that both free thiamin and TPP over-producing lines exhibited altered carbon partitioning in the seeds. Seed phenotype analysis revealed that transgenic lines accumulated more oil and carbohydrate, but less protein than the control lines (Table 1 and 2). To date, the attempts to increase seed oil content in plants were mostly focused on altering the expression level of the genes and/or transcription factors involved in the oil biosynthetic pathway (Zou et al., 1997; Jako et al., 2001; Vigeolas et al., 2007; Kim et al., 2014; Van Erp et al., 2014, Kim et al., 2019; Guo et al., 2020). It has been shown that the fatty acid biosynthesis pathway is highly dependent upon NADH and NADPH supplies (Geer et al., 1979; Slabas and Fawcett, 1992). There are some controversial reports regarding the sources of reductant supply for oil biosynthesis. Kang and Rawsthorne (1996) suggested that NADPH generated by oxidative PPP possibly provides reducing power for fatty acid biosynthesis. On the other hand, Rawsthorne (2002) reported that NADPH generated in the chloroplasts might be a potential source for oil biosynthesis. Using a quantitative metabolic flux model, Schwender et al. (2003) suggested that glycolysis and the oxidative PPP can provide the reductant required for fatty acid biosynthesis. Nevertheless, the sources of reducing molecules have not been identified yet (Schwender et al., 2003). In our study the significant increase in seed oil content observed in thiamin and TPP over-producing lines might be due to the higher levels of reductants produced by thiamin cofactor-dependent enzymes such as TK, PDH, and α-KGDH (Jordan, 2003; Nosaka, 2006), as their transcript levels were also significantly increased in high-thiamin lines (Fig. 8). Additionally, the oil biosynthetic pathway is dependent on acetyl-CoA supply which is required for growing the acyl chain in the pathway (Slabas and Fawcett, 1992). The significant increase of *PDH* gene expression indicated in this study suggest a possible way for elevating the acetyl-CoA level needed for producing more oil in transgenic lines. To support our findings, it is noteworthy to consider that increased oil content caused by high levels of thiamin is not limited to the higher plants. In algae, Higgins et al. (2016) showed that a significant increase in lipid content can be achieved by the addition of HMP, free thiamin, and/or TPP to the algae medium.

Fatty acid composition analysis also showed some differences between transgenic and control seeds. Indeed, 3-gene overexpressing lines contained higher levels of palmitic, stearic, and Oleic acids but lower levels of linolenic acid compared to the wild type seeds (Table 3). These results are consistent with the results reported by Higgins et al. (2018) in which they showed that by the application of HMP and thiamin in the medium supplemented with glucose as organic substrate, algae cells reserved higher levels of palmitic and Oleic acids but the lower levels of linolenic acid.

Our results revealed that carbon flux shifted from protein to oil biosynthesis in *Arabidopsis* and *C. sativa* transgenic seeds (Table 1 and 2). It has been shown that overexpression of *DIACYLGLYCEROL ACYL TRANSFERASE1* (*DGAT1*) and *WRINKLED1* (*WRI1*) along with the RNAi suppression of the lipase *SUGAR-DEPENDENT1* (*SDP1*) could increase the oil content of the *Arabidopsis* seeds; however, a significant decrease in seed protein content was also observed (Van Erp et al., 2014). Additionally, using mutational analysis, Focks and Benning (1998) reported that *WRI1* has an indirect effect on fatty acid biosynthesis. They suggested that *wri1* mutants have aborted the conversion of glucose into fatty acid precursors, and sucrose and hexoses concentrations were increased in developing *wri1* mutant seeds. Maisonneuve et al. (2010) also reported that increased seed oil content in transgenic *Arabidopsis* overexpressing rapeseed lysophosphatidic acid acyltransferase (LPAAT), an important enzyme needed to synthesize the phosphatidic acid required for storage lipid biosynthesis in developing seeds, could be at the expense of seed storage proteins. Furthermore, Kanai et al. (2016) showed the negative correlation between oil and protein contents during *Arabidopsis* seed development. On the other hand, because of the participation of N and S atoms in thiamin structure (Jurgenson et al., 2009), the substantial decrease in seed protein level in high-thiamin and TPP transgenic lines can be partly attributed to the fact that these lines use protein as N and S sources to produce pyrimidine and thiazole rings of the thiamin. Together, these results indicate that plants use protein and sugar as the carbon source for oil biosynthesis (Focks and Benning, 1998) and also demonstrate the competition between the metabolic pathways for the substrates (Schwender and Hay, 2012; Van Erp et al., 2014; Kanai et al., 2016).

Increased seed oil content in both the thiamin and TPP over-producing lines might also be a consequence of increased carbohydrate levels generated by the up-regulation of *TK* gene due to high level of thiamin cofactor (Fig. 8). TK enzyme has a significant role in carbon flux in the Calvin cycle (Haake et al., 1999; Raines et al., 2000) and elevated sugar content in transgenic seeds might be the consequence of the increased regeneration level of ribulose-1, 5-bisphosphate, which is a substrate for the Rubisco, by TK enzyme. This is consistent with the fact that in tobacco plants, antisense *TK* transformants displayed reduced photosynthesis and sugar content due to the inhibition of ribulose-1, 5-bisphosphate regeneration (Henkes et al., 2001). These results suggest that metabolic pathways which rely on plastid TK are highly sensitive to slight changes in plastid *TK* expression (Henkes et al., 2001).

Oil, storage proteins, and carbohydrates in the form of starch are the major constituent in the seeds of flowering plants (Baud et al., 2002; Wang et al., 2007). Seed reserves provide initial carbon and energy source for growing seedlings (Bradbeer, 2013). During morphogenesis, oil in the form of TAG is accumulated in *Arabidopsis* embryos to be used for seed germination and seedling establishment (Mansfield and Briarty, 1992; Eastmond et al., 2000). Analysis of hypocotyl growth in control and thiamin over-producing seeds revealed that transgenic seeds produced a longer hypocotyl than the control seeds (Fig. 9). These results can be attributed to the fact that higher levels of thiamin in transgenic lines provide more TPP, which is essential for boosting carbohydrate, oil, and protein metabolism required for hypocotyl growth when photosynthesis is blocked by complete darkness.

Seed mass was another trait that was analyzed in this study and a significant increase in seed weight of the thiamin over-producing genotypes was observed compared to the control lines (Fig.7B). Consistent with our results, Jako et al. (2001) reported a remarkable increase in seed weight ranging from 40% to 100% by the overexpression of *DGAT* gene in *Arabidopsis*. In addition, overexpression of rapeseed *LPAAT* gene, led to a boost in oil content and seed mass in *Arabidopsis* (Maisonneuve et al., 2010). Moreover, expression of yeast LPAAT isozymes in *Arabidopsis* and rapeseed remarkably enhanced seed oil and seed weight (Zou et al., 1997). Kanai et al. (2016) also reported that overexpression of *WRI1* gene in *Arabidopsis* resulted in the production of larger seeds, and the weight of the seeds was up to 1.7-fold greater than those of control. On the other hand, Zhang et al. (2005) showed that silencing of *DGAT1* gene in tobacco reduced both mature seed oil content and average seed weight, while an increase in protein and soluble sugar content of the mature transgenic seeds was observed. These results indicate that the larger seed mass of the high-thiamine lines is in part due to altered carbon partitioning between the seed primary storage compounds.

## MATERIALS and METHODS

### Plant materials and growth conditions

Seeds of wild type *Arabidopsis* plants (ecotype Columbia) and *Camelina sativa* (cv. Celine) were used in this study. Wild type *Arabidopsis* seeds were grown on full strength MS (Murashige and Skoog, 1962) agar plates at 21°C under 100 µmol m^-2^ s^-1^ constant light for 10 days. They were then transplanted to the soil for growth under 130 µmol m^-2^ s^-1^ light with 16 h light/8 h dark period in a growth chamber. Twenty five-day-old plants were used for floral dip transformation by *Agrobacterium tumefaciens* (Clough and Bent, 1998). *C. sativa* seeds were spread directly on the soil until they grow and form unopened buds under 300 µmol m^-2^ s^-1^ light with 16 h light/8 h dark period in a growth chamber. Then, floral dip transformation was carried out as described by Liu et al. (2012).

### Creation of DNA constructs for the overexpression of thiamin biosynthesis genes

To overexpress the *AtThi4* (At5G54770) and *AtThiC* (At2G29630) genes in wild type *Arabidopsis* and *C. sativa* seed, two gene expression cassettes were made (Fig. 2A) in which the *AtThi4* and *AtThiC* open reading frames (ORFs) were cloned into the NotI site of pMS4 and pKMS2 under the control of *Brassica* napin (Ellerström et al., 1996) and oleosin (Keddie et al., 1994) promoters, respectively. The pMS4 and pKMS2 vectors were then excised in their AscI sites to remove the fragments of napin promoter::*AtThi4*::glycinin 3′UTR and oleosin promoter::*AtThiC*::oleosin terminater, respectively. These two expression cassettes were then sub-cloned into the AscI and MluI sites of the RS3GseedDSredMSC binary vector. The *AtThiE* (At1G22940) ORF was cloned directly into the XmaI site of RS3GseedDSredMSC vector under the control of soybean (*Glycine max*) glycinin promoter (Iida et al., 1995). The *Agrobacterium tumefaciens* strain GV3101 was then transformed using binary vector harboring these 3 genes. Transgenic *Arabidopsis* and *C. sativa* seeds (T_1_) were produced using *Agrobacterium*-mediated transformation by the floral dip method (Clough and Bent, 1998). The T_1_ seeds were then selfed to produce the T_2_ seeds. To identify the homozygous lines, T_2_ plants were grown and segregation analysis was performed on the siliques of *Arabidopsis* and also on *C. sativa* pods containing the T_3_ seeds against the *DSRed* gene using fluorescent microscope. In parallel, homozygous plants harboring the empty vector control were produced to be used for further analysis.

For the overexpression of *AtTPK1* (At1g02880) gene, *AtTPK1* ORF was cloned into the XmaI site of the RS3GseedDSredMSC binary vector to have soybean glycinin promoter:: *AtTPK1*::glycinin terminator. PCR was performed to amplify the *AtTPK1* gene with the glycinin promoter and terminator. This fragment was next sub-cloned into the NotI site of the pART27 binary vector (Fig. 2B). The homozygous transgenic and empty vector control lines were produced as described for 3-gene overexpressing lines, except we used kanamycin to select the transgenic lines.

### Seed thiamin content analysis using HPLC

Dry seeds (∼5 mg) from wild type, empty vector control, and transgenic lines were ground in 300 µL of 2% (w/v) trichloroacetic acid (TCA). Samples were incubated at 95 °C for 30 min and kept on ice for 2 min, and then centrifuged at 14,000g for 5 min. The supernatant was centrifuged using Nanosep Centrifugal Filter Device (0.2 mm) columns (Pall Life Sciences) for 3 minutes. Free thiamin and TPP in 2% TCA were then converted into thiochrome using cyanogen bromide (Kim et al., 1998). Thiochrome peaks were identified by HP 1046A Fluorescence Detector (Agilent Technologies, Palo Alto, CA) at 370 nm excitation and 430 nm emission using HP 1100 HPLC (Agilent Technologies, Palo Alto, CA) equipped with a Capcell Pak NH_2_ column (150 mm × 4.6 mm i.d., Shiseido, Tokyo). A 4:6 (v/v) solution of 90 mM potassium phosphate buffer, pH 8.4 and acetonitrile was used as the mobile phase (Ajjawi et al., 2007b).

### RNA isolation and real-time quantitative RT-PCR (qRT-PCR)

Total RNA was isolated from 9-day-old developing *Arabidopsis* siliques (Van Erp et al., 2014) using the Spectrum^TM^ Plant Total RNA Kit (Sigma-Aldrich). The isolated total RNA was then treated with DNase I (Amplification Grade, Sigma-Aldrich) according to manufacturer’s protocol. First strand cDNA was synthesized utilizing 2 µg of total RNA and iScript^TM^ cDNA Synthesis Kit (Bio-Rad). A fraction (0.75 μg) of the cDNA was used as the template in 10 μl reaction mixture per well in real-time PCR. TaqMan probes were purchased for *Thi4*, *ThiC*, *ThiE*, *TPK1*, sytosolic *TK* (AT245290), chloroplastic *TK* (AT3G60750), cytosolic *PDH* (AT1G24180), chloroplastic *PDH* (AT2G34590), cytosolic and mitochondrial *α-KGDH* (AT3G55410), mitochondrial *α-KGDH* (AT5G65750), chloroplastic deoxy-xylulose phosphate synthase, *DXPS3*, (AT5G11380), and chloroplastic *DXPS2* (AT4G15560) (Assay ID. No. At02243600_g1, At02252013_g1, At02305767_g1, At02333096_gH, At02263110_g1, At02197798_g1, At02306409_g1, At02216989_g1, At02189484_g1, At02283104_g1, At02273275_g1, and At02221416_g1, respectively) from Applied Biosystems (Life Technologies, Carlsbad, CA). The *Arabidopsis eEF-1 α* gene (Assay ID. No. At02337969_g1) tagged with VIC dye (Life Technologies, Carlsbad, CA) was added in each well as an internal control. PCR thermal cycling conditions used for amplification was 95 °C for 2 min followed by 39 cycles of 95 °C for 15 sec., 60 °C for 1 min. Gene expression levels were quantified by CFX96 Real-Time System (Bio-Rad). Data was analyzed using 2^-ΔΔCT^ method (Tunc-Ozdemir et al., 2009).

### Salt and paraquat stress assays

In all stress experiments the seeds were surface sterilized and kept at 4 °C for 2 days in complete darkness and then germinated at 22 °C under 100 µmol m^-2^ S^-1^ constant light in growth chamber. Germination assays were performed with at least five technical replications, each containing 50 seeds per line, and repeated at least three times. For seed germination assay, the seeds were grown on full strength MS agar plates without adding thiamin, and containing the different concentrations of NaCl and paraquat (methyl viologen) for 5 days and the number of germinated seeds were counted each day. As a morphological marker, visible penetration of the radicles into the seed coat was considered for seed germination (Kim et al., 2008). These plates were also kept in the growth chamber for 4 more days to evaluate the number of viable seedlings for each genotype.

For root growth assay, the seedlings (5 seeds per line for *A. thaliana* and 4 seeds per line for *C. sativa* with at least 10 technical replications) were grown vertically on full strength MS agar plates without adding thiamin, supplemented with various concentrations of NaCl and paraquat. The increase in root length was measured each day for 9 days. The RGR was then calculated as described by Lupini et al. 2010.

### Seed lipid analysis using GC

In *A. thaliana*, seed oil analysis was performed according to Li et al. 2006. Two mg of seed was measured out and aliquoted to glass tubes. A 300 µl of the mixture of toluene and triheptadecanoin (total of 96 µg triheptadecanoin per sample) was added to each sample, then 25 µl of butylated hydroxytoluene solution (0.2% butylated hydroxytoluene in MeOH), and finally 1 ml of sulfuric acid (5% H_2_SO_4_ in MeOH). The samples were incubated at 95 °C for 1.5 h and then allowed to cool to room temperature for 30 minutes. A 1.5 ml of 0.9% NaCl (w/v) was added to each sample. Finally, a 2 ml hexane extraction was performed on each sample twice by vortexing to mix and then centrifugation for phase separation. The top supernatant layer of each extraction was removed and pooled into a fresh glass tube and then dried down using a stream of N_2_ (g). Samples were then re-suspended in 400 µl of the hexane and transferred to a GC vial. Oxidative air was purged from each GC vial using a gentle stream of N2 (g). Samples were run through a DB-23 column (Agilent Technologies, Santa Clara, CA) using 2 µl injections in a 25:1 split ratio. Injector and FID temperatures set to 260 °C. Prior to starting the run, the column was conditioned at 240 °C for 2 hours. For each sample, the oven temperature started at 50 °C for each run and was ramped up to 150 °C at 25°C/minute and then held at 150 °C for 3 minutes. The oven was then ramped to 240 °C at 10 °C/minute, and then the temperature was held at 240 °C for 5 minutes. For oil data analysis, the raw data for each chromatogram was transferred to an excel format. The data was processed by labeling each major FAME peak using the external standards as a guide. Analysis was performed by normalizing the area under each peak to the area of the C17 internal standard peak to produce a normalized peak area. The normalized peak area was multiplied by the mass of internal standard added to each sample (0.096 mg) for an amount of lipid per peak, which was then divided by the total sample weight (2 mg) and multiplied by 100% to produce an oil content relative to dry weight. The peaks for each sample were summed to produce a total lipid content for that sample. The data for each sample was analyzed for an average value, standard error, and a 95% confidence interval.

To measure the oil percentage in *C. sativa* seeds, Benchtop NMR (Bruker NMR mq20) was used according to manufacturer’s instruction. Canola oil was used as a standard.

### Seed protein and carbohydrate assay

Seed protein content was measured as described by Focks and Benning (1998). Briefly, 20 to 50 seeds were weighed out and homogenized in 250 µl of acetone in a 1.5-ml tube. Following centrifugation at 16,000*g*, the supernatant was discarded and the vacuum-dried pellet was resuspended in 250 µl of extraction buffer containing 50 mM Tris-HCl, pH 8.0, 250 mM NaCl, 1 mM EDTA, and 1% (w/v) SDS. Following a 2 h incubation at 25 °C, the homogenate was centrifuged at 16,000*g* for 5 min and 5 µl of the supernatant was used for protein measurement in a microplate format utilizing the Lowry DC protein assay kit and γ-globulin for generating standard curve (Bio-Rad).

For quantification of total carbohydrate, 50 seeds were homogenized in 500 µL of 80% (v/v) ethanol in a 1.5-ml tube and incubated at 70 °C for 90 min (Focks and Benning, 1998). The total sugar was measured using the phenol-sulfuric acid method in microplate format and glucose was used to generate standard curve (Masuko et al., 2005). Soluble sugars were analyzed according to Focks and Benning (1998).

### Hypocotyl length assay

To perform the hypocotyl length assay, surface sterilized seeds were plated on agar plates containing half strength MS salt without adding thiamin and sucrose. The seeds of each genotype were plated in different positions to minimize the plate position effects. The edge of the plates was then wrapped with 3M Micropore surgical tape. Following wrapping the plates in aluminum foil, they were kept in 4 °C for 2 days in complete darkness to break the seed dormancy. The aluminum foil was then removed and the plates were placed in 22 °C under 100 µmol m^-2^ S^-1^ constant light for 24 h. After 24 h, the plates re-wrapped in aluminum foil individually and placed vertically in 22 °C. The hypocotyl length was measured for 8 and 6 days for *A. thaliana* and *C. sativa*, respectively.

### Statistical analysis

Data analysis was performed using Student’s *t* test (Suzuki et al., 2008) to show the significance of differences between data sets.

## Acknowledgement

We would like to thank Professor Edgar Cahoon’s lab at the University of Nebraska-Lincoln for providing pMS4, pKMS2, and RS3GseedDSredMSC vectors.

